# SUPERmerge: ChIP-seq coverage island analysis algorithm for broad histone marks

**DOI:** 10.1101/121897

**Authors:** Bohdan B. Khomtchouk, Derek Van Booven, Claes Wahlestedt

## Abstract

SUPERmerge is a ChIP-seq read pileup analysis and annotation algorithm for investigating alignment (BAM) files of diffuse histone modification ChIP-seq datasets with broad chromatin domains at a single base pair resolution level. SUPERmerge allows flexible regulation of a variety of read pileup parameters, thereby revealing how read islands aggregate into areas of coverage across the genome and what annotation features they map to within individual biological replicates. SUPERmerge is especially useful for investigating low sample size ChIP-seq experiments in which epigenetic histone modifications (e.g., H3K9me1, H3K27me3) result in inherently broad peaks with a diffuse range of signal enrichment spanning multiple consecutive genomic loci and annotated features.

## Introduction

Standard ChIP-seq wet and dry lab workflows are very well characterized, and their respective tools and protocols frequently and consistently employed [2, 16, 22]. One hallmark shared in common by all ChIP-seq computational workflows is the process of peak calling, whereby differential enrichment of reads is statistically detected across the genome in case-control samples.

It is known that epigenetic histone modifications that have inherently broad peaks with a diffuse range of signal enrichment (e.g., H3K9me1, H3K27me3) differ significantly from narrow peaks that exhibit a compact and localized enrichment pattern (e.g., H3K4me3, H3K9ac) [15]. Likewise, transcription factors that exhibit punctate protein-DNA enrichment patterns resulting in narrow and well-defined peaks differ significantly from the broad peaks frequently detected in diffuse chromatin mark data. Due to these differences in peak topology, various specialized peak calling algorithms have been developed to handle both transcription factor ChIP-seq and histone modification ChIP-seq data analysis [6, 11–14, 20, 24, 28, 29, 31, 32].

Despite these computational advances in identifying putatively enriched genomic regions, peak calling software generally relies on pooling the respective BAM files of each biological replicate within each group (in cases and controls) prior to performing the peak calling step. This is done in order to merge multiple sorted alignment (BAM) files within each group, thereby reliably boosting read count and amplifying the downstream differential enrichment signal obtained during the peak calling process. While pooling certainly achieves the aforementioned goal, it may inadvertently lead to erroneous results due to one or two over-representative samples whose read counts swamp the overall signal. This is especially relevant to experiments with low sample sizes and yields (e.g., brain tissues [3,19]), where a single biological replicate might drive the overall signal of the (case or control) group. As such, it is strongly recommended to pre-process data with quality control software to ensure that the sequencing phase was successful [27]. A popular tool designed for this purpose is FastQC [1], which can check for biases and other sequencing anomalies, including duplication rate and per-base GC content and sequence quality score. A complementary tool, MGA [10], can check for contamination, un-alignable sequence, and presence of sequencing adapter dimers [27]. In addition, the irreproducible discovery rate (IDR) can be used to measure general consistency between replicates in high-throughput experiments [17] and is made available as an R package [18]. Another popular alternative is the R/Bioconductor package ChIPQC [7], which computes a number of quality metrics relating to the alignments, including duplication rates, mapping quality filtering rates, distribution of pileups (and the associated SSD measure), and an estimated fragment size based on maximizing a relative cross-coverage score [26].

Despite the abundance of quality control and peak calling software available to-date, there has not yet been a tool for quantitatively examining read pileup and annotation information at the level of a single BAM file. This would be useful for identifying read pileup outliers across select individual (un-pooled) samples, as well as investigating the latent impact of broad coverage islands whose summit position, length, and other dimensions are unlikely to be registered as statistically significant (i.e., false-negatives) by existing peak calling software. Since peak caller statistical significance often precludes many genomic loci (and their respective genes) from further consideration, SUPERmerge offers the analyst complete freedom in specifying coverage depth parameters and broadness dimensionality (see Methods section) – a paradigm-shifting idea that we propose for annotating genomic features and planning subsequent follow-up analyses. As such, we recommend using SUPERmerge for exploratory data analysis purposes (not peak calling), e.g., when investigating annotations of extremely broad histone modification coverage islands that span multiple consecutive genomic loci and annotated features.

## Methods

SUPERmerge is a cross-platform command-line program written in the C programming language that takes as input a single BAM file and produces as output a variety of tabulated flat file results (see software manual), as well as graphical data visualization output (see Results section) generated by the R programming language [23]. SUPERmerge source code and compilation instructions are made publicly available at: https://github.com/Bohdan-Khomtchouk/SUPERmerge. A precompiled binary executable and software manual are available for immediate download at: https://sourceforge.net/projects/supermerge/files/SUPERmerge.zip/download.

SUPERmerge is designed to be used as part of a genomic workflow (Figure 1) prior to, or in conjunction with, the process of peak calling (see Results and Discussion sections for rationale). The program is composed of three primary parameters that are specified as user flags:

- −*d*: depth
- −*i*: interval grouping factor
- −*g*: GTF annotation

**Figure 1.**
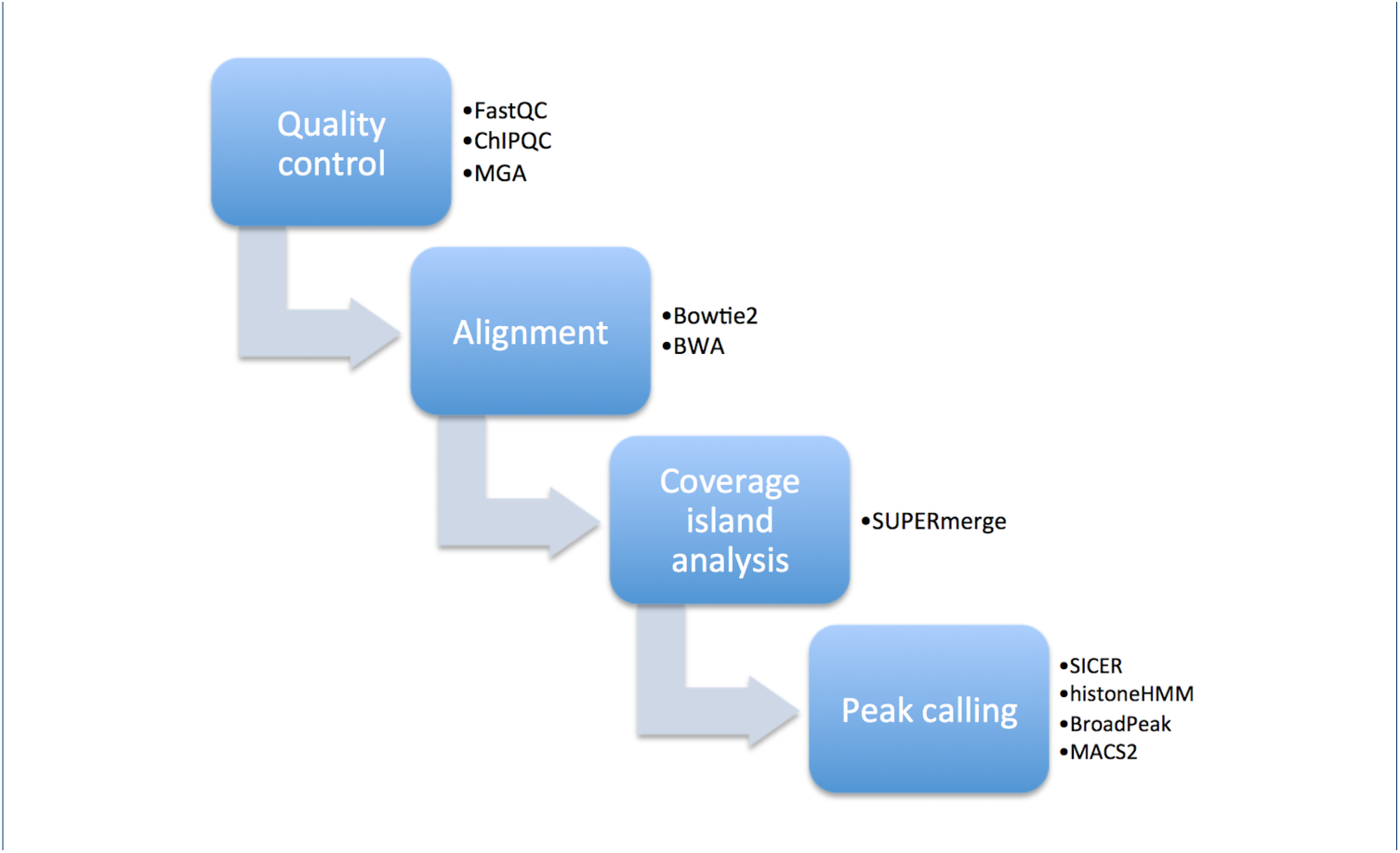
Standard ChIP-seq workflow. Following the quality control and read alignment steps, SUPERmerge is designed to be run as an exploratory data analysis tool either prior to or in tandem with the peak calling step. Details of the SUPERmerge algorithm are presented in Methods.

Depth represents read pileup at a given genomic location (e.g., “−*d* 20” returns all single base pair positions in a BAM file harboring at least 20 or more reads). SUPERmerge leverages the mpileup function in the SAMtools software suite [?] to calculate depth. The interval grouping factor (−*i*) represents a genomic distance (in bp) between any two consecutive positions of sufficient depth (−*d*), for which any two neighboring base pair positions are considered to be part of the same genomic interval if they are separated by no more than this distance (−*i*). In practice, this translates to merging any base pair positions that are of sufficient depth *−d* into one contiguous interval of coverage, as long as the gap between these genomic positions is no greater than −*i*. We refer to these contiguous regions of coverage as “coverage islands”, since there exists no base pair position within a −*i* radius of the island (in either the 3-prime or 5-prime end) that is of the sufficient depth −*d*. However, within any given coverage island, there will always be a certain percentage of base pairs that do not meet the sufficient depth, simply by virtue of the fact that any two consecutive regions of sufficient depth are merged into a continuous interval whenever there is a gap of less than −*i* bp between them. The frequency of occurrence of coverage islands whose base pairs meet a certain percentage of sufficient depth per island is counted and graphed in Figure 2A (see Results section for details).

**Figure 2.**
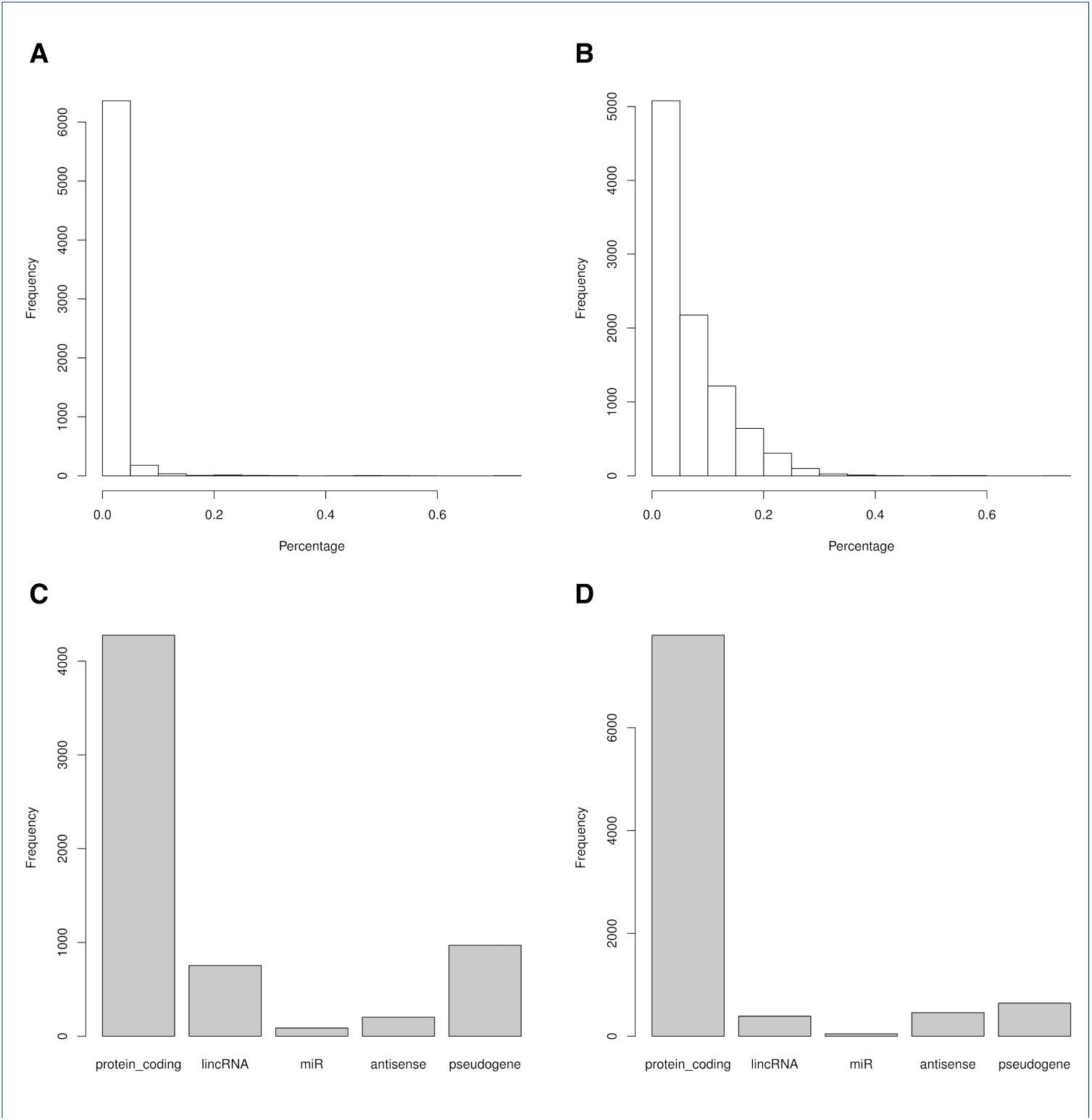
Percentage & annotation graphs for two biological replicates from a sample histone modification ChIP-seq experiment. Panels A and C depict SUPERmerge runs for replicate 1, while Panels B and D depict results for replicate 2 for an experiment where sample size *n* = 2. Please see Results for details.

As such, these two thresholds (−*d* and −*i*) constitute the cutoff parameters by which a user can explore how reads assemble into coverage islands at various levels of exploratory data analysis. Coverage islands are defined as contiguous genomic regions with a certain percentage of read coverage above a set threshold depth and a distance of at least −*i* bp from their nearest neighboring island. Once these coverage islands are calculated, the −*g* flag specifies a gene transfer format (GTF) file that annotates them with their respective genomic features (see Results section).

An example of a full command issued from the command-line is:

~~~
./supermerge -d 20 -i 500 -g gencodeV19annotation.gtf file.bam
~~~

This command would be repeated by the user across all BAM files in the sample pool, for any desired combinations of −*d* and −*i*.

## Results

SUPERmerge allows users to analyze read pileup distribution across individual alignment (BAM) files of biological replicates, returning a variety of graphical and tabulated output results specific to the parameter space described in Methods. There are four graphs that are produced for each set of replicate BAM files:

- Annotation graph: a bar chart of all coverage islands annotated to GTF features (protein_coding, lincRNA, miR, antisense, and pseudogene)
- Island diameter graph: a histogram of coverage island length vs. frequency of occurrence
- Library size graph: a histogram of coverage island area (total sum of reads per island) vs. frequency of occurrence
- Percentage graph: a histogram showcasing the percentage of bases within an individual coverage island that meet the sufficient depth (−*d*) after merging (−*i*) vs. frequency of occurrence

Figure 2 shows a representative histone modification ChIP-seq sample run produced for the H3K4me3 mark in the human GM12878 cell type downloaded from the ENCODE ChIP-seq Experiment Matrix [5, 9]. This data is composed of two biological replicates, both of which are run under the same set of parameters for their respective flags (−*d*, −*i*, and −*g*):

~~~
./supermerge -d 20 -i 500 -g gencodeV19annotation.gtf
    wgEncodeBroadHistoneGm12878H3k4me3StdAlnRep1.bam
./supermerge -d 20 -i 500 -g gencodeV19annotation.gtf
    wgEncodeBroadHistoneGm12878H3k04me3StdAlnRep2.bam
~~~

Figures 2A and 2B depict the number of coverage islands in each biological replicate that meet a sufficient read depth (e.g., “frequency *x* at percentage *y*” means that there exists a total of *x* coverage islands wherein only *y* percent of the base pairs in each respective island meet the required depth −*d*). As explained in Methods, this is due to the nature of the −*i* parameter in merging contiguous genomic regions of sufficient depth (−*d*) that are sufficiently close to each other in distance (within −*i* bp).

Despite an approximately similar set of annotations in the two biological replicates (Figures 2C and 2D), the distribution of coverage islands that meet the sufficient depth is drastically different (Figures 2A and 2B). In fact, most coverage islands in replicate A have less than 10% of base pairs that meet the sufficient depth (−*d* = 20), suggesting a significantly more diffuse signal enrichment profile relative to replicate B. Likewise, coverage islands in replicate A exhibit a significantly lower variance in length (measured from end-to-end of each island) across the BAM file relative to replicate B (Figures 3A and 3B). Furthermore, the library size graph of replicate A indicates that there are many more outliers of coverage islands that contain extremely large quantities of reads per island, despite having a lower median value than replicate B (Figures 3C and 3D). This suggests that replicate A is dominated by a large number of oversized coverage islands that, taken in consideration with Figure 2A, contain an abundance of shallow reads that do not meet the sufficient depth. It is precisely this abundance of reads failing to meet the sufficient depth that gives rise to such a high number of coverage island outliers with huge library sizes (i.e., total sum of reads per island). Taken together, this suggests that replicate A contains an elevated count of scattered contiguous regions of sufficient depth that are closely separated (within 500 bp of each other), suggesting a more fractured alignment pattern than replicate B. By studying the percentage graph of replicate A at progressively diminishing values of −*i* (Figure 4A-F), it becomes clear that coverage islands in this replicate sample are separated by approximately 100 bp from each other on average. Comprehensive analyses of percentage, annotation, island diameter, and library size graphs for both biological replicates are provided as supplementary materials for download here: https://github.com/Bohdan-Khomtchouk/SUPERmerge/tree/master/output/interval_figures.

**Figure 3.**
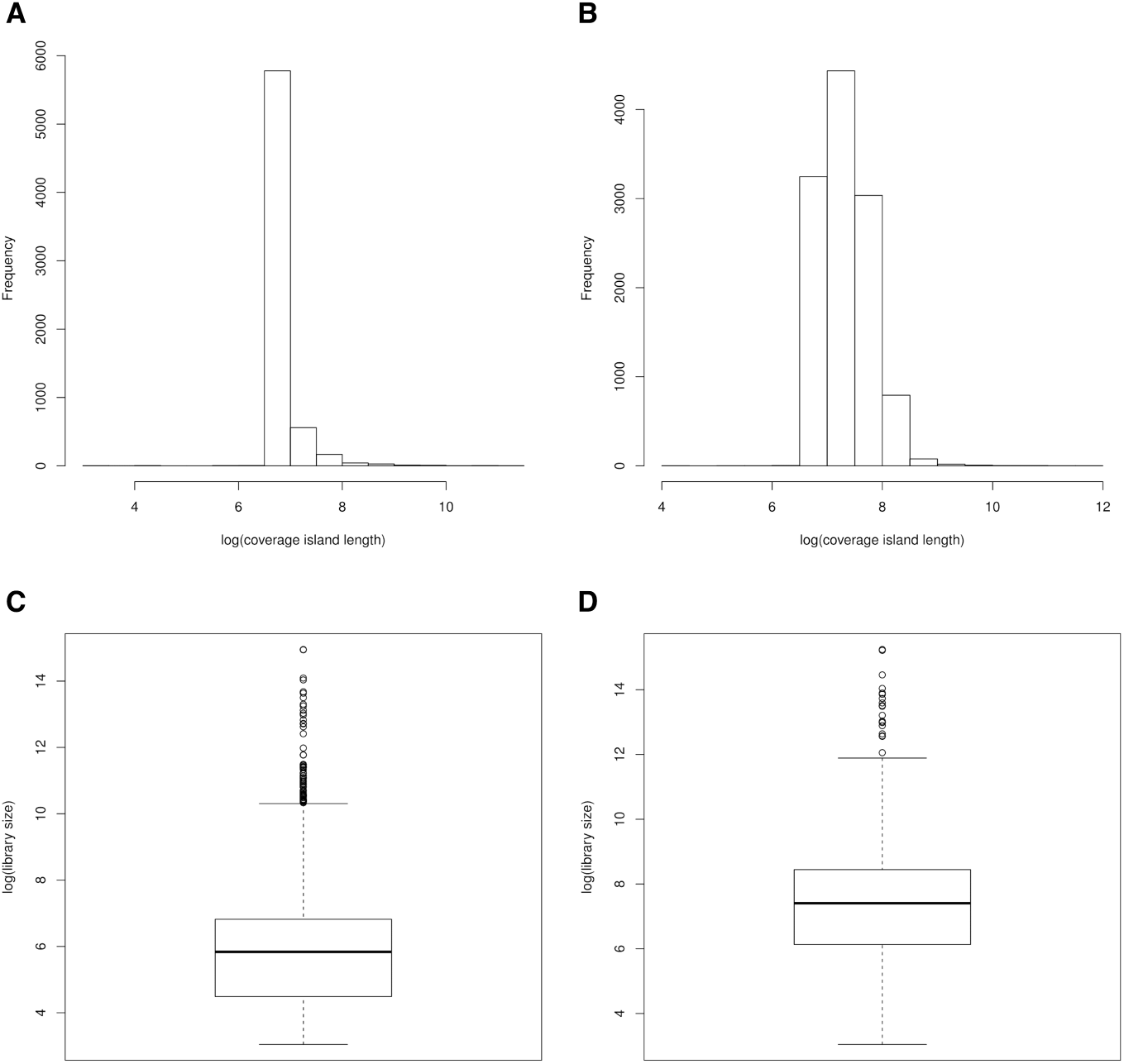
Island diameter & library size graphs for two biological replicates from a sample histone modification ChIP-seq experiment. Panels A and C depict SUPERmerge runs for replicate 1, while Panels B and D depict results for replicate 2 for an experiment where sample size *n* = 2. Please see Results for details.

**Figure 4.**
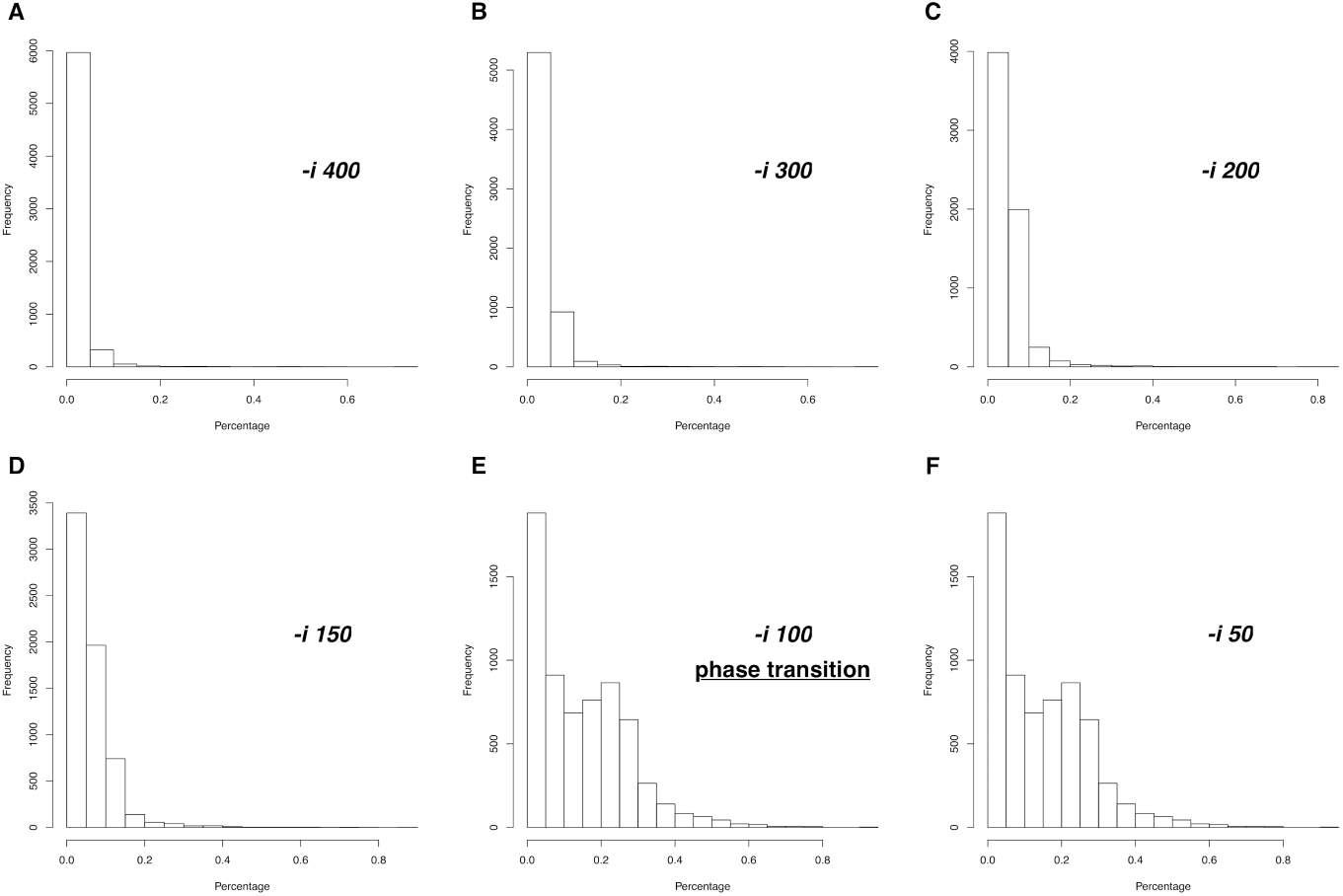
Phase transition in coverage island percentage graph at constant depth cutoff. Panels A through F represent gradually decreasing values of −*i* at a constant value of read depth (−*d* = 20). In the transition from −*i* = 150 to −*i* = 100, a phase transition is seen in the abundance of coverage islands possessing higher percentages of reads that meet the sufficient depth relative to the total reads within each island. Practically no visible change is seen from −*i* = 100 to −*i* = 50, suggesting that coverage islands in this biological replicate of this histone modification ChIP-seq dataset are, on average, surrounded by approximately 100 bp of read desert (where “read desert” is defined by the user as < 20 reads per base pair position for approximately 100 consecutive base pairs in both the 3-prime and 5-prime directions).

## Discussion

The analysis of coverage islands in the histone modification ChIP-seq experiment presented as a case-study in Results highlights the underlying freedom of the user in testing different sets of input parameters in the SUPERmerge algorithm for exploratory data analysis purposes. Since many histone modification ChIP-seq experiments are composed of only a few biological replicates per group, this approach is likely to prove useful in making critical downstream analysis decisions, such as whether or not and how to pool or remove replicates during the subsequent peak calling and differential enrichment steps. Considerations involving the specific biology of various chromatin marks will also likely inform the results. For instance, the H3K4me3 study presented above suggests a relatively narrow domain of separation between coverage islands for this specific histone mark, which is in line with its known biological role as the methylation state associated with transcriptional start sites of actively transcribed genes [4].

Since large-scale ChIP-seq analyses are known to be sensitive to the quality of input samples, this has often necessitated the need to filter poor quality data automatically [21]. However, in experiments with naturally low sample sizes and yields (e.g., brain tissue [3,19]), this is a risky and wasteful procedure. Even though larger numbers of biological replicates are known to establish better concordance and extracted signal [30], this is clearly not always possible. Seeing as the sample size issue will likely continue to impact the ChIP-seq field in many areas of tissue research (e.g., brain), SUPERmerge has been designed to mediate and navigate the aforementioned conflicts by allowing users to explore within-group characteristics of individual alignment (BAM) files, thereby highlighting differences in read alignment topology across single biological replicates of the same group and aiding the user in planning subsequent analysis decisions. All in all, this approach is designed to complement and inform the many peak caller analysis approaches currently in widespread use by the scientific community.

## Conclusions

We have designed a novel bioinformatics algorithm implemented as a cross-platform command-line program, SUPERmerge, for the analysis and annotation of coverage islands within individual read alignment (BAM) files of histone modification ChIP-seq datasets harboring broad chromatin domains. Coverage islands are defined as contiguous genomic regions of read pileup that are isolated from their nearest neighbors by some genomic distance (in bp) on both the 3-prime and 5-prime ends. By examining coverage island formations at a single base pair resolution across individual biological replicates, SUPERmerge is designed to be used as an exploratory data analysis tool that complements existing peak calling and quality control algorithms to inform users when to pool or remove samples from analysis based on the consistency and concordance of biological replicates to each other.

## Author information

Center for Therapeutic Innovation and Department of Psychiatry and Behavioral Sciences, University of Miami Miller School of Medicine, 1120 NW 14th St., Miami, FL, USA 33136. Correspondence.

## Author‘s contributions

BBK conceived the study and wrote the paper. BBK and DV wrote the code. CW participated in the management of the source code and its coordination. All authors read and approved the final manuscript.

## Acknowledgements

BBK dedicates this work to the memory of his uncle, Taras Khomchuk. BBK wishes to acknowledge the financial support of the United States Department of Defense (DoD) through the National Defense Science and Engineering Graduate Fellowship (NDSEG) Program: this research was conducted with Government support under and awarded by DoD, Army Research Office (ARO), National Defense Science and Engineering Graduate (NDSEG) Fellowship, 32 CFR 168a.

## Abbreviations used

bp: base pairs
ChIP-seq: chromatin immunoprecipitation followed by massively parallel DNA sequencing
IDR: irreproducible discovery rate
QC: quality control
TF: transcription factor

